# Residence time analysis on cerebral aneurysms treated with coils using planar-laser-induced fluorescence and computational fluid dynamics

**DOI:** 10.1101/2022.10.18.512786

**Authors:** Julia Romero Bhathal, Syed Faisal, Michael Levitt, Christian Geindreau, Alberto Aliseda

## Abstract

Biofidelic numerical models have been developed such as the coil-resolved model to study hemodynamics in the treated aneurysm. In this model, the geometry of the coils is recreated from high-resolution tomography scans of a phantom aneurysm treated with coils. However, this model hasn’t been validated. The purpose of this work is to validate the coil-resolved model. To achieve this, we used the planar-laser induced fluorescence technique on phantom aneurysm treated with coils and measured the residence time and the evolution of rhodamine concentration during the washout. We run passive scalar simulations with the coil-resolved model and measured the evolution of concentration over time. The comparison of the numerical and the experimental results shows that the coil-resolved model reproduces the hemodynamics of the experimental setup. Therefore it can be used as a reference to study hemodynamics in the treated aneurysm or to validate porous media models developed for treatment outcomes prediction.

## 1. Introduction

Cerebral aneurysms (CA) are often treated with endovascular coiling. This technique is preferred due to its non-invasiveness and shorter post-operative recovery time for patients, compared to craniotomy surgery. It consists of deploying platinum coils inside the aneurysmal sac, slowing down blood flow into the aneurysm and inducing thrombus formation. The aneurysm is then isolated from the stresses associated with blood flow, reducing the risk of wall rupture and subarachnoid hemorrhage [1, 2]. The number of coils placed inside the aneurysm depends on the aneurysm anatomy, with the objective to fill at least 25-30% of the aneurysm volume. Thrombosis, however, is not always complete [3]. Recanalization can lead to high risk of continuous growth of the aneurysm and potential rupture. There are factors of the coil deployment inside the aneurysmal sac, like porosity, that have been investigated to define the probability of success of the treatment [4]. However, these parameters do not provide a global predictive metric for stable thrombus formation after treatment, and there is currently no agreement on how to predict the success or failure at the time of treatment.

Research on this problem has focused on numerical modeling of blood flow in the cerebral aneurysm. Hemodynamics has been strongly connected with aneurysm growth and rupture [5]. Determining which hemodynamics factors are involved in thrombus formation, is the first step towards prediction of treatment outcomes [6]. Numerical models of the coiled aneurysm, such as developed by [7], accurately represents the physiology, and can be used to determine parameters linked with recanalization. This coil-resolved model reproduces patient-specific boundary conditions, and uses the exact geometry of the coils, which cannot be obtained from a clinical scan. It represents an improvement on current numerical modeling that has been attempted to use in translational settings, but hasn’t been validated experimentally (*in vivo* or *in vitro*). Studying hemodynamics *in vivo* in coiled aneurysms presents a significant challenge and has not been documented in the literature. Indeed, the coils produce artifacts on most imaging modalities and, due to its opacity, reduce the field of view and accuracy when visualizing blood flow *in vivo*.

Thrombus formation is linked with low blood velocity and stagnation in the aneurysmal sac. Residence time in the aneurysm can be a useful parameter to determine the future development of thrombosis. It is expected that a high residence time will lead to a stable thrombus formation, and better treatment outcomes [8].

Previous studies have used planar laser-induced fluorescence (PLIF) to measure concentration of species over time in in *vitro* experiments [9, 10]. This technique could allow to visualize the washout out in a aneurysm treated with coils, using a fluorence dye and following the evolution of its concentration in the aneurysm over time, and to measure residence time in the treated aneurysm.

The aim of the present work is to determine residence times for a small cohort of patients, both numerically through computational fluid dynamics and experimentally through planar laser-induced fluorescence (PLIF). The second objective is to evaluate the consistency of the results between the two methods, validating the coil-resolved computational model.

## 2. Methods

### 2.1. Model creation

Six patients were enrolled in this study at the University of Washington’s Harborview Medical Center in Seattle, WA, USA. They all presented cerebral aneurysms and were treated endovascularly with coils (Stryker Endovascular, Kalamazoo, Michigan, USA). All coils had the same diameter, 240 — 250 μm, with lengths between 2 — 30 cm. Each patient received a different treatment (number of coils and lengths) based on the anatomy and the neurosurgeon’s judgment. For each patient, the parent vessel and the aneurysm were imaged using three-dimensional rotational angiography. After image segmentation, a numerical 3D model of the aneurysm and the parent vessel was created for each patient. A silicone model, at 1:1 scale, was built in acrylonitrile butadiene styrene with 3D printing technology. The silicone model was then cast in a clear polyester resin (PDMA, Clear-Lite; TAP plastics, San Leandro, California, USA) [11]. This transparent model was treated with the same set of coils as the patient; the same surgeon performing the surgery in the patient placed the coils in the models, in the same order and with the same specifications as those received by the patient. The aneurysm volumes can be found in Table 1: patients E and F present small volumes, C and D medium volumes and G and H large volumes. Patients were selected based on variance in aneurysm volume to gain better understanding of how the volume of the aneurysm impacts residence time.

**Table 1:** Volume of the aneurysm, mean porosity (*ϕ_m_*), pore volume for Patients C to H

| Patient | Aneurysm volume (mm<sup>3</sup>) | $\phi_m$ | $V_{pore}$ (mm<sup>3</sup>) |
| --- | --- | --- | --- |
| C | 93.8 | 0.772 | 72.4 |
| D | 90.0 | 0.715 | 64.3 |
| E | 32.9 | 0.678 | 22.3 |
| F | 38.2 | 0.720 | 27.5 |
| G | 413.3 | 0.795 | 328.5 |
| H | 279.9 | 0.788 | 220.6 |

Coils cannot be accurately imaged in a clinical scan, therefore, the coils placed in the silicone model were imaged with high-resolution tomography, avoiding beam hardening artifacts, at the European Synchrotron Radiation Facility (beamline ID19, ESRF, http://www.esrf.eu). The six models were imaged at a resolution of approximately 13 μm. After the treated models were scanned, the coils were segmented from the reconstructed 3D images using ImageJ [12], and the surface of the coils was extracted as a mesh (stl format). The process from the patients’ medical images to coil surface Synchrotron images is shown in Figure 1.

**Figure 1:**
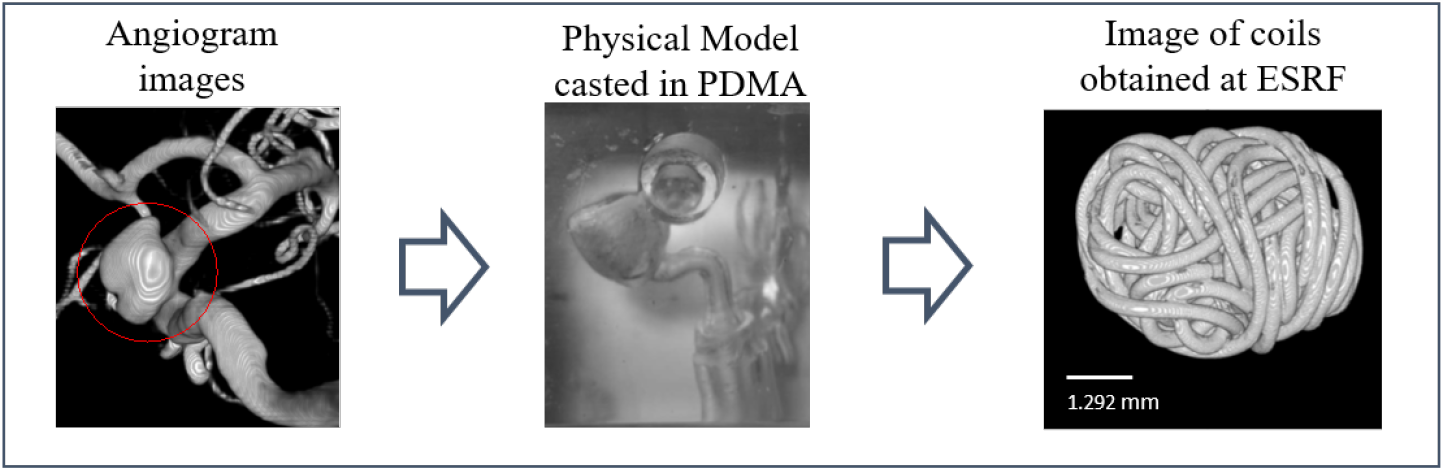
Experimental pipeline: From angiographic image (left), to 3D-printed model (middle) in which coils are placed, to synchrotron scans of the coil mass (right)

The coils’ surface geometry from the synchrotron images was used to create a numerical model of the aneurysm and parent vessel treated with coils for computational fluid dynamics (CFD) analysis. The coil surface was positioned in the numerical 3D model of the aneurysm and parent vessel (from the CT scans). The process involved extracting the centerline of the parent vessel in the numerical 3D model and the 3D synchrotron images. The centerlines were extracted using Vascular Modeling Toolkit software (http://www.vmtk.org). The iterative closest point method [13] allowed us to find the rotation and translation matrices between the centerlines of the parent vessels (numerical and synchrotron models). These same matrices were applied to the geometry of the coils, resulting in a numerical coil-resolved model that reproduced exactly the geometry of the experimental model (treated with coils) placed in the same reference frame as the pre-treatment scan of the parent vessel lumen. The coil-resolved models obtained in this study are shown in Figure 2.

**Figure 2:**
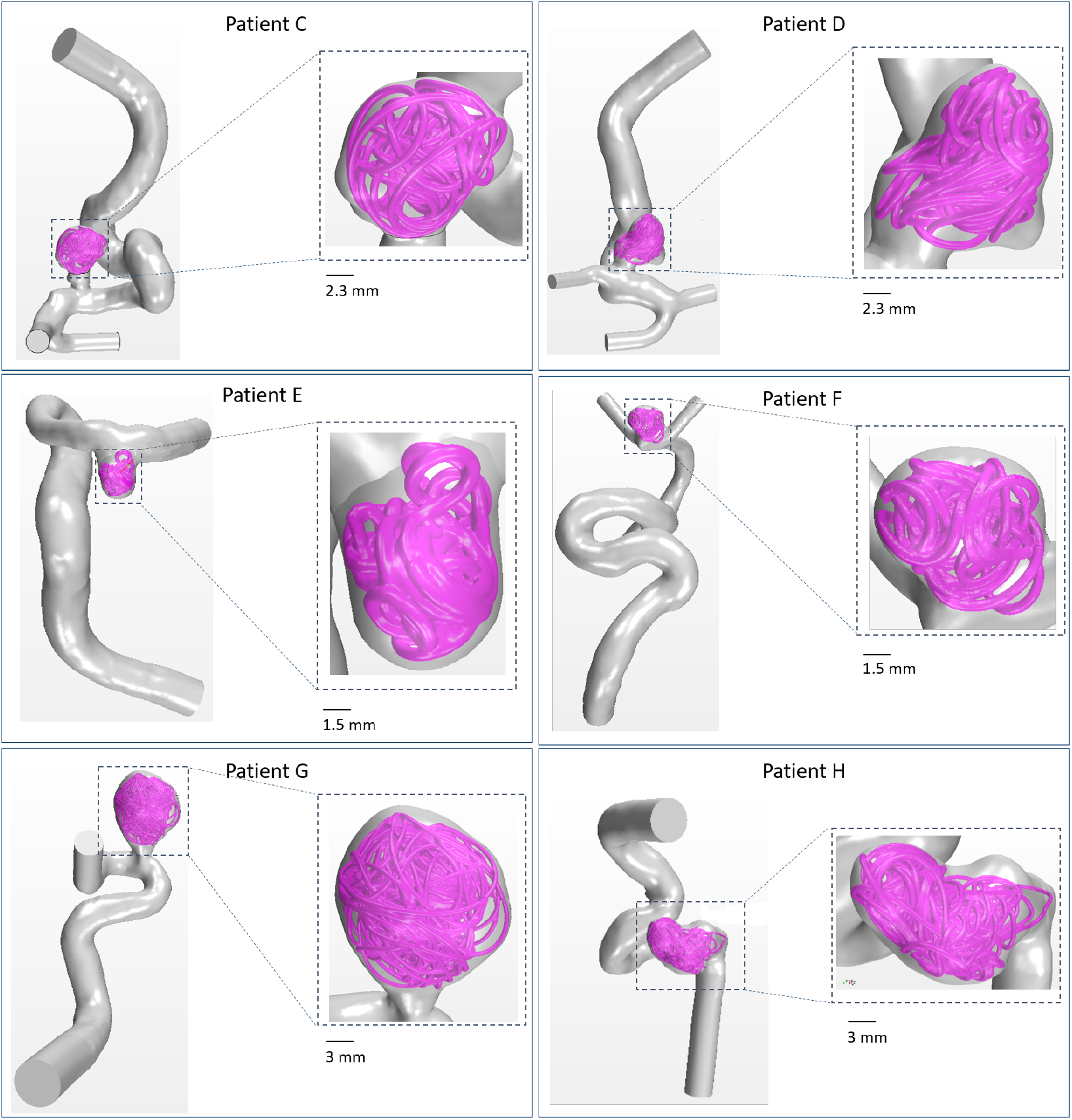
3D models of the patients after coils reposition.

### 2.2. Planar-laser-induced fluorescence

#### 2.2.1. Experimental set up

For the experiment, the phantom artery was connected to a flow loop that reproduced the parent vessel hemodynamics (see Figure 3). The flow loop consisted of a reservoir placed above the experiment, to avoid air entrainment in the flow loop. A pulsatile pump (Harvard Apparatus, Boston, MA, USA), mimicking the waveform in the aneurysm parent artery, pumped the working fluid through the system. A resistance and capacitance were placed downstream from the pump to reproduce the effect of the cardiovascular system. The pump settings, resistance, and capacitance were adjusted so the volumetric flow rate at the parent vessel in the imaging test section was *Q_mean_* = 200 mL/min and *Q_systole_* = 2*Q_mean_* [14]. The heart rate set for the pulsatile pump was 55 BPM, with a ratio of duration for systole/diastole of 40/60. The flow rate was measured with an ultrasonic flow meter (Titan, Dorset, UK). The model was placed downstream from the flow meter. The tubes and fittings were adapted for each model depending on the patient’s anatomy (one or multiple outlets). However, the rest of the boundary conditions were the same for all patients. The working fluid (a transparent solution of 47.5% water, 35.8% glycerol, and 16.7% NaCl) was chosen to match the viscosity of blood (3.8 cp) and the index of refraction of the silicone, avoiding any refraction artifacts in the images [15].

**Figure 3:**
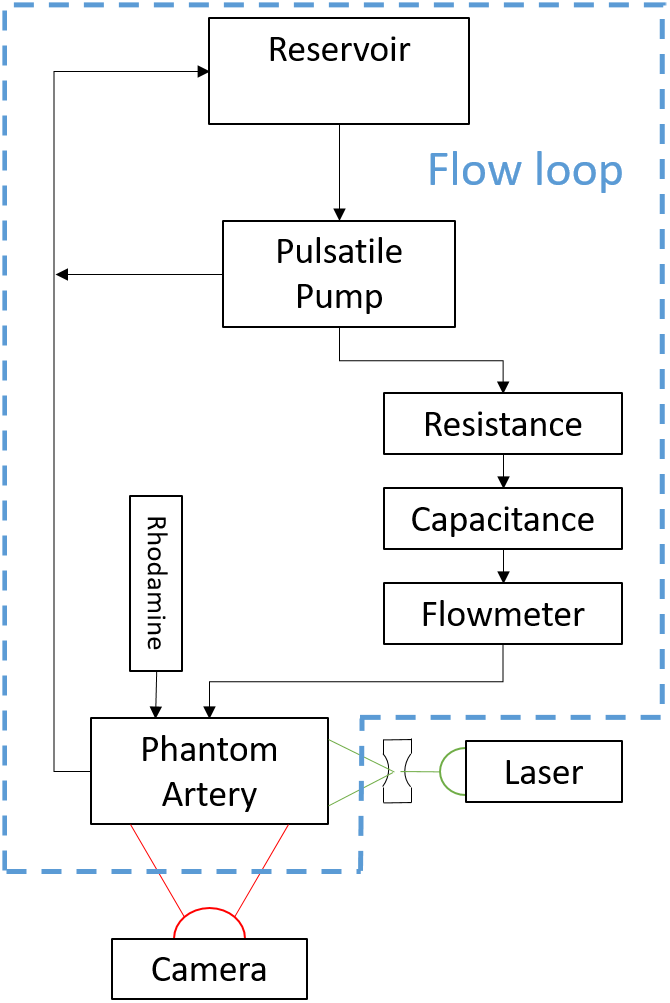
Schematics of the optical set up and the flow loop

A solution of rhodamine B in the working fluid, at a concentration of 5 ppm, was created and injected in the aneurysmal sac prior to the start of the experiment. To visualize the initial fluid washout from the aneurysm sac, measuring the residence time of blood in the aneurysmal sac, the rhodamine B solution that filled the aneurysm was illuminated by a laser at the frequency of excitation of rhodamine B (Coherent Genesis STM CX Laser at 532 nm) and visualized with a high-speed camera. The laser light was shaped into a plane, focusing the beam into a thin beamline and expanding into a plane (1.1 mm thickness) using cylindrical and spherical lenses. The high-speed camera focal axis was perpendicular to the laser plane. Further details of the imaging technique are presented in the next section.

The experiment was primed by injecting the rhodamine B solution into the aneurysmal sac, slowly through a needle that was placed in the sac through the walls of the model (not the parent vessel) avoiding disturbances in the aneurysmal sac, confirmed by imaging prior to the start of flow. Once the aneurysm was filled, the pump was turned on and the physiological flow in the parent vessel was started, with some of it entering the aneurysmal sac and pushing out some of the dyed fluid that filled the sac initially. The full process of washout of the rhodamine solution was recorded by high-speed camera.

Figure 3 illustrates the flow loop and the optical setup, where the arrows illustrate the flow direction. Figure 4 shows the phantom artery for Patient G, with the arrows representing inflow and outflow, the aneurysm treated with coils, and the needle through which the rhodamine was injected.

**Figure 4:**
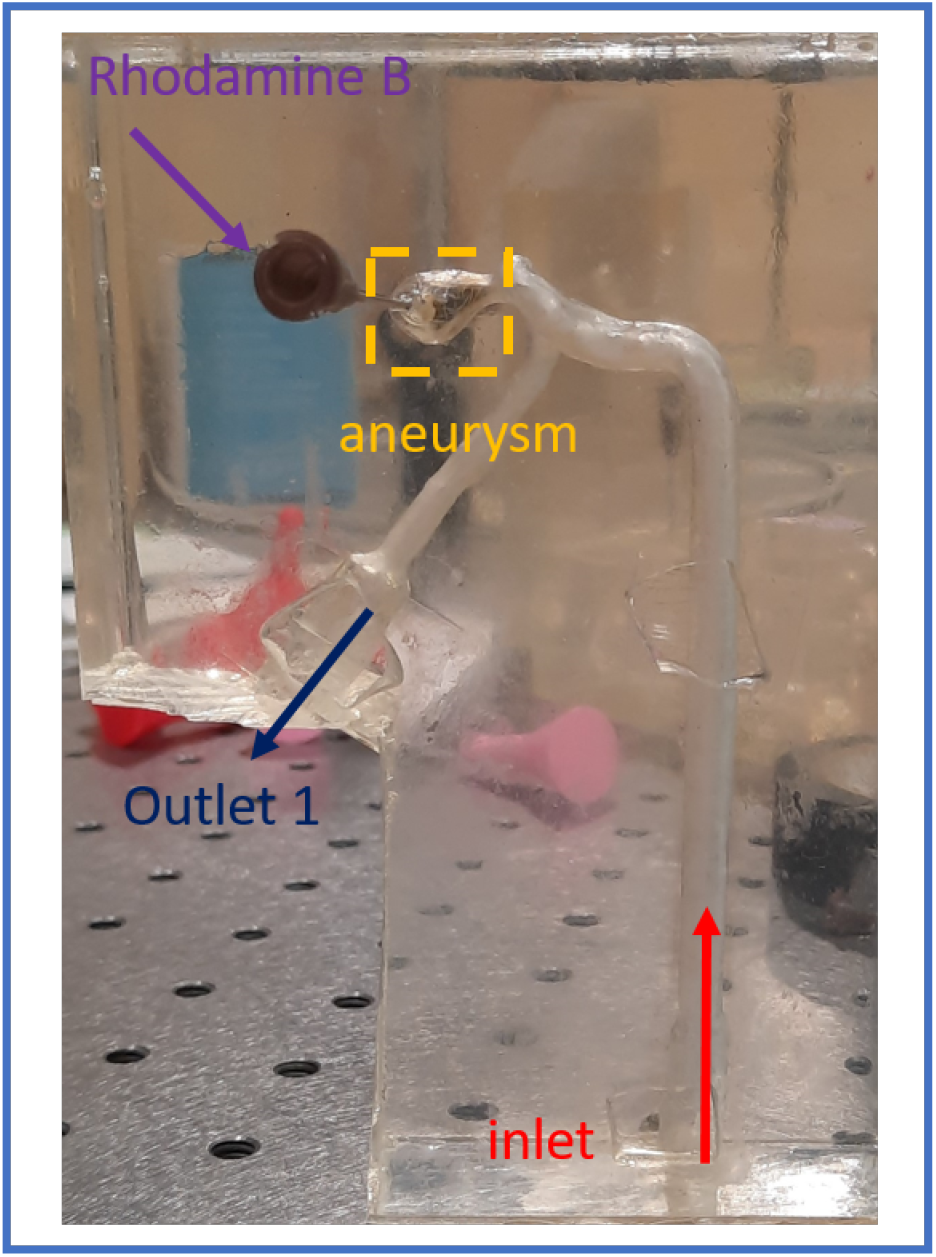
Silicone model patient G.

#### 2.2.2. Image acquisition and processing

A high-speed camera (Phantom v711, Vision Research, Trenton, NJ) captured the fluorescent light emitted by the rhodamine B dye after being excited by the laser light. A band-pass filter was placed between the camera and the artery phantom to remove the reflections of incident light. The rate of image acquisition and length of time captured in the experiments depended on the size of the aneurysm, varying between 70 seconds at 100 fps (for the largest aneurysm) and 14 seconds at 500 fps (for the smallest aneurysm).

The images recorded were treated with ImageJ [12]. First, the last image, for which the washout was completed, was subtracted from all the images to remove the signal from the coils and other spurious reflections that had gone through the band-pass filter. Then, the image intensity was thresholded to segment the sections were rhodamine B was present and those clear of fluorescent dye. This image-processing operation also limited the artifacts created by the presence of coils. The aneurysm was defined as the convex envelope of the coils. The rhodamine B that could have leaked into the parent vessel during injection was excluded from the analysis by the aneurysm envelop segmentation, before calculating the concentration of rhodamine. The analysis of the residence time was done using Matlab (Mathworks Inc., Natick, MA). The overall volume of rhodamine in the image sequence was calculated as the sum of the total pixel intensity values within the image, normalizing the overall intensity integrated value in the first image to 100%. The residence time was defined as the time at which the volume of rhodamine in the aneurysm first decayed by 99%, that is when it reached 1% in the normalized intensity integration. The process for image analysis is outlined in Figure 5.

**Figure 5:**
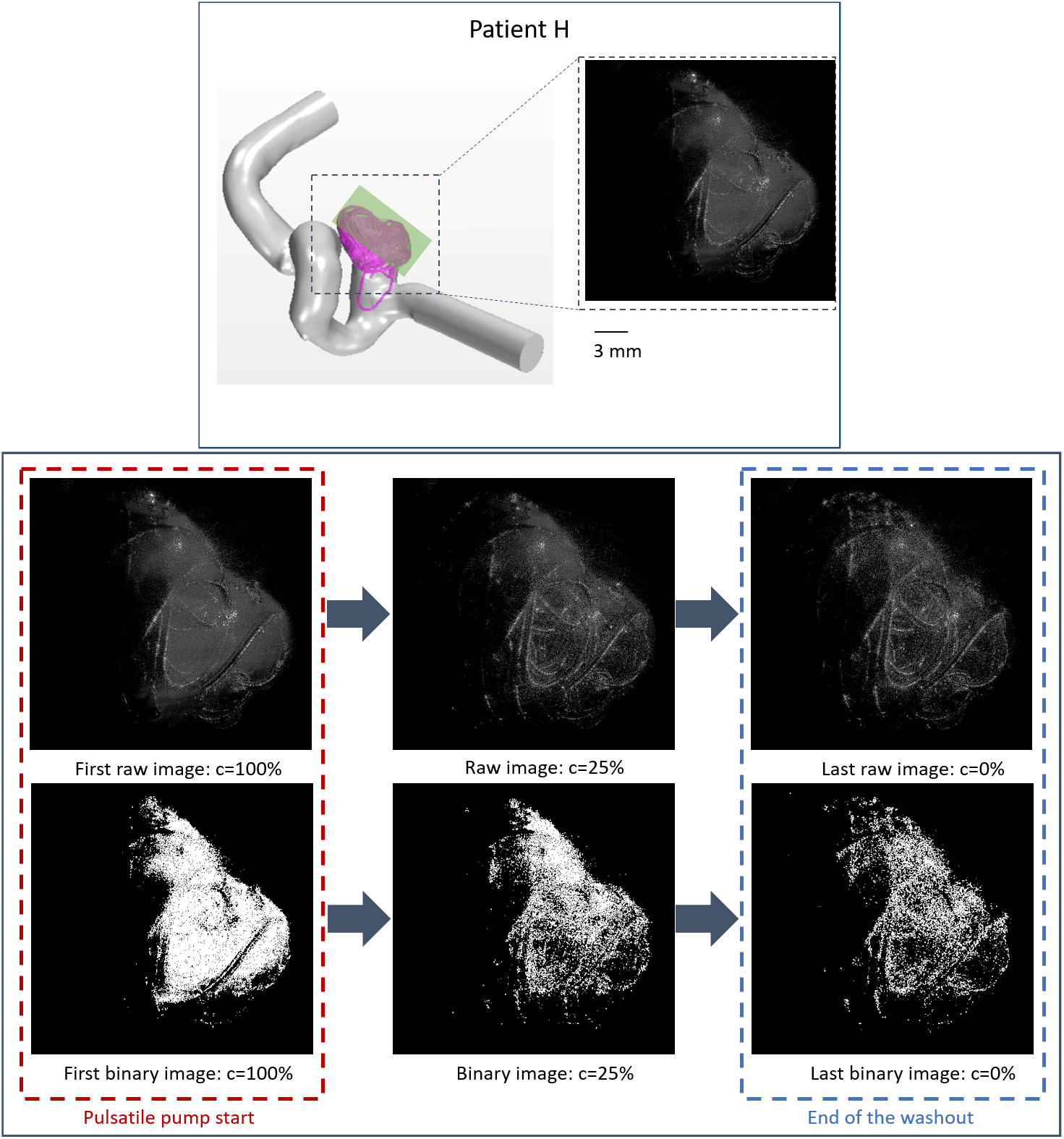
Patient H image treatment: on the top, the 3D model of Patient H with the green plane that is equal to the images obtained through PLIF. On the bottom left the images (raw and binary) before starting the pulsatile pump, when the aneurysm is filled with rhodamine B (c=100%). On the middle, at c=25%, and on the right (raw and binary) at the end of washout (c=0%).

#### 2.3. Computational Fluid Dynamics

The experimental setup described above was used to validate the coil-resolved computational model. From each of the six patients, residence time was calculated using the finite volume flow simulation. The reconstructed geometry detailed in previous sections (see Figure 2), was meshed with a tetrahedral mesh. The element size of the parent vessel was 200 μm and the element size at the surface of the coil was 20–40 μm. This process was performed using StarCCM+ [7].

Blood was modeled as an incompressible Newtonian fluid, with viscosity equal to 0.0035 Pa.s and density equal to 1050 kg/m^3^ [16]. The ssoftware used for this analysis was Fluent (ANSYS, Release 17.1; ANSYS, Canonsburg, Pennsylvania, USA). In the following section, we describe the method for the numerical simulation of the physics and the boundary conditions used in the simulations.

##### 2.3.1. Physics

The Navier-Stokes equations were integrated numerically to simulate the blood flow in the aneurysms and the convective-diffusive transport of the dye:

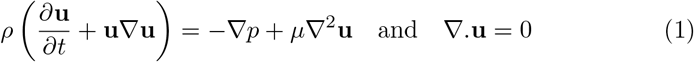

where *ρ* is the blood density and *μ* the blood viscosity, **u** and *p* are the fluid velocity and the pressure respectively. To study residence time in the aneurysm treated with coils, the results of the experimental model were compared with the numerical model. In the experimental study, the aneurysm was filled with rhodamine, and to calculate the residence time, the concentration of rhodamine was calculated during the washout with the transport equations for incompressible flow:

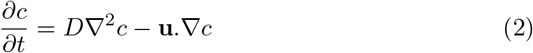

where *c* is the concentration of a passive scalar, and *D* its diffusivity. In Fluent (ANSYS, Release 17.1; ANSYS, Canonsburg, Pennsylvania, USA), this equation is solved to study the evolution of rhodamine concentration in the aneurysm over time. At *t* = 0s (begining of the washout), *c* = 1 in the aneurysm and *c* = 0 in the parent vessel. The diffusivity of rhodamine in water at 21.5°C is *D* = 3.6 × 10^-6^ cm^2^.s^-1^, and this value is used for solving equation (2) in the CFD simulations.

##### 2.3.2. Boundary conditions and residence analysis

The boundary conditions in the CFD model were patient-specific. At the inlet, a pulsatile Womersley velocity flow was imposed, reconstructed from measurements performed *in vivo* by the surgeon using a dual-sensor Doppler guidewire (ComboWire and ComboMap; Volcano Corp, San Diego, California, USA). Those measurements were used to reproduce a pressure condition for the patients in the outlets [17]. For patients with only one outlet, a zero-pressure condition was imposed in the outlet. For patients with multiple outlets, Resistance-Capacitance conditions were used. The arterial wall and the surface of the coils were considered rigid with a non-slip boundary condition.

In the experimental study, the boundary conditions were fixed for all the patients. However, in the CFD model, the boundary conditions varied from one patient to another. The comparison between CFD and experimental results was done by calculating the evolution of the rhodamine B concentration versus the inlet volume flow rate in the experimental study, and the evolution of concentration versus the inlet volume flow rate imposed for each patient in the numerical study.

Simulations ran for five cardiac cycles. The first two cycles were discarded as they are influenced by the transient effects on the simulation. The rhodamine concentration was set to 1 in the aneurysmal sac at the beginning of the third cycle and to 0 in the parent vessel and the inlet. The washout was analyzed during the remaining three cardiac cycles. For some cases, due to the size of the aneurysm, the washout was not complete at the end of the third cardiac cycle. However, due to the long computational time required to run the simulations, it was decided to only run three cardiac cycles for each patient, to calculate the trend of the evolution of the concentration of rhodamine.

### 3. Results

#### 3.1. PLIF results

The results of the PLIF study are presented in Figure 6. This figure shows the evolution of the rhodamine B concentration over time for each patient. Five runs were recorded for each patient and the three runs presenting the most consistent initial conditions for the concentration at the beginning of the experimental measurements were analyzed.

**Figure 6:**
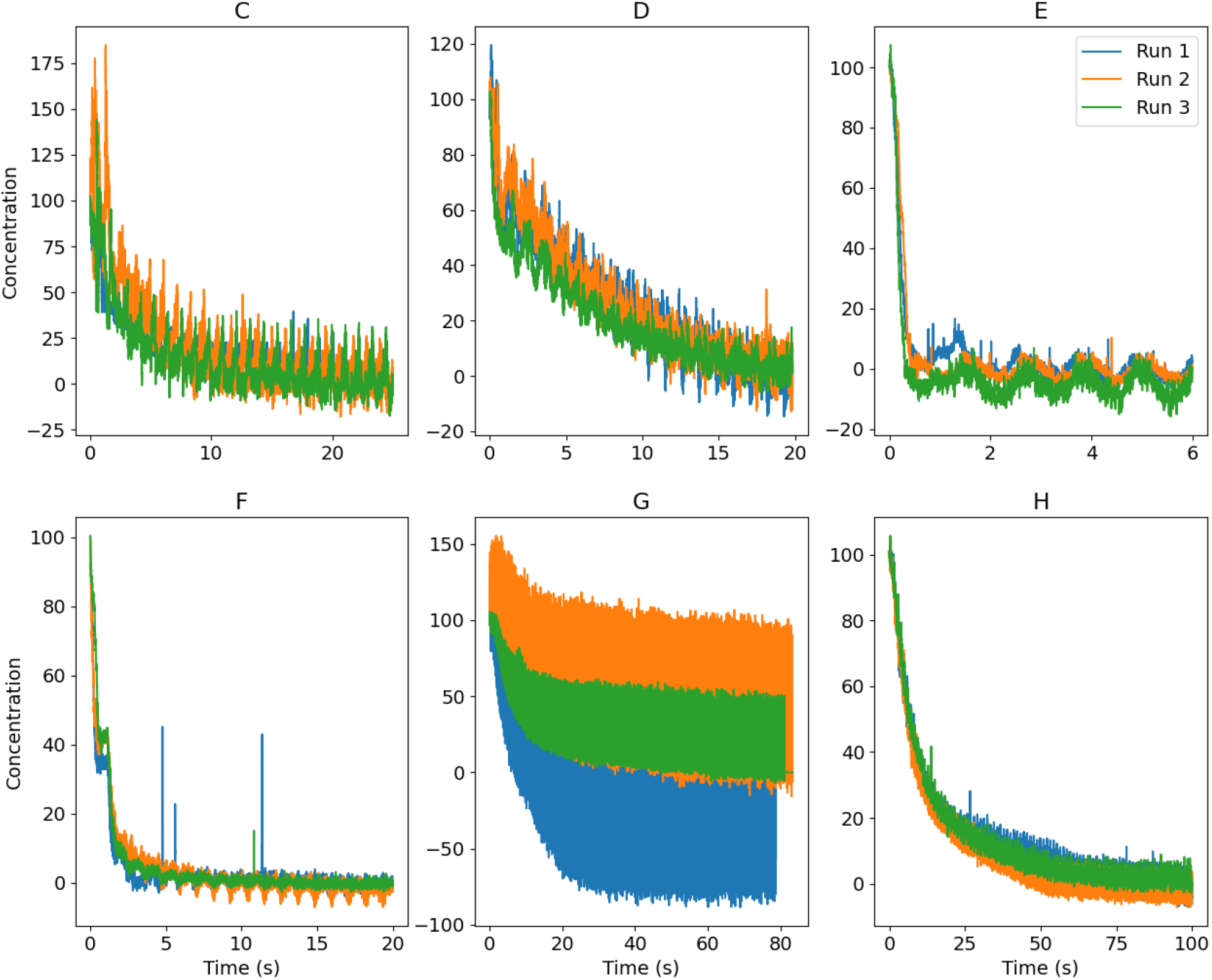
Evolution of the concentration during time for each patient over three run for the PLIF analysis.

The quality of the data was impacted by the size and the location of the aneurysm and the laser plane chosen. The impact of the size can be easily identified when analyzing the data in Figure 6: Patients E and F present the clearest signal while being the smaller aneurysms. It is more difficult to visualize the dye fluorescence in larger aneurysms because it is harder for the laser plane to penetrate the larger coil mass and illuminate the complete aneurysm volume. This can be visualized in Figure 7. The green plane shows the area that was visualized in the camera and where the concentration was calculated. For small aneurysm patients as E and F, the totality of the aneurysm was illuminated, whereas for bigger aneurysms only a part of the aneurysm filled with rhodamine was visualized.

**Figure 7:**
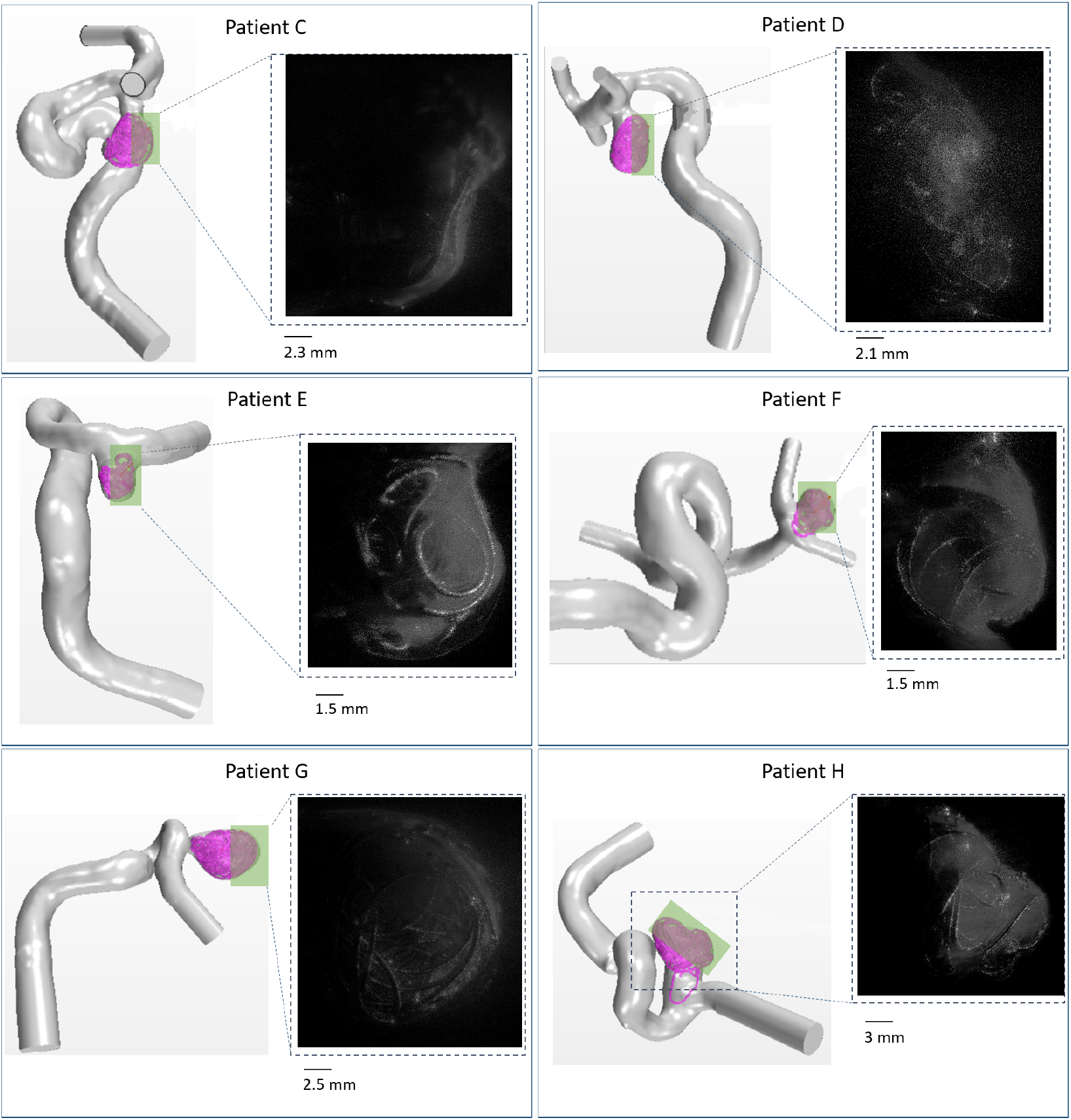
3D numerical model vs image obtained through PLIF technique for each patient. In each numerical model, the green plane shows region observed in the images obtained through PLIF techniques, which correspond to the area illumanated with the laser. The PLIF image used is when the rhodamine concentration in the aneruysm is at 100%. The 3D models are positionned as they were in front of the camera.

The quality of the model (clarity of the silicone walls) also impacts the quality of the data but is minor when looking at the impact of the location of the aneurysm in the anatomy. Placing the model in the setup is highly difficult due to the limitations of the plane size and to the positioning of the needle used to inject the rhodamine. The model was set up to avoid that the parent vessel would intercept the laser plane, or in the camera view while avoiding visualizing the needle to avoid image artifacts due to the metallic composition of the needle. However, for some models as for Patient D, the laser light goes through the parent vessel resulting in poor image quality as it was impossible to place in a different position (see Figure 7D).

The laser plane is chosen to influence the analysis of the evolution of rhodamine B concentration in the aneurysm over time. For Patient G, the evolution of rhodamine concentration is studied in the area located on the opposite side of the neck (see Figure 7G). That region corresponds to the lower velocities in the aneurysm filled with coils, and the coils create artifacts due to the reflection. This creates a noisy signal which explains Figure 6G. The laser plane is conditioned by the two previous factors: the size of the aneurysm and the anatomy of the parent vessel and aneurysm.

The cardiac cycle (systole and diastole) also has an impact on the quality of the data. When analyzing Patient E (Figure 6E), even when the washout is complete there is still some variation in the concentration. Indeed, when the laser reaches the coils it created artifacts due to reflection. The coils move during the cardiac cycle which leads to a variation in the light intensity perceived with the camera during the cardiac cycle, creating this variation in the pixel values even when the washout is complete. Patients C and D present more variation in the results than patients E and F because they were recorded for a longer time as their aneurysm volume was larger. Nevertheless, those patients present a clear evolution of rhodamine concentration over time.

Figure 8 presents the evolution of the rhodamine concentration in the aneurysm over time normalized by the aneurysm volume for each patient. The concentration is the mean of the concentration over the three runs for each patient. Patient G was excluded from this comparison due to the impact of the noise in the results visualization. The results show that all the patient present the same trend: the concentration of rhodamine decrease rapidly until at least 50% of the rhodamine is cleared away. Then the concentration decreases at a slower pace until complete washout. This phenomenon is amplified for the smaller aneurysms. Indeed Patient E presents a decrease of the rhodamine concentration very fast until its concentration is around 5% in the aneurysm, and then slowly decreases until full washout.

**Figure 8:**
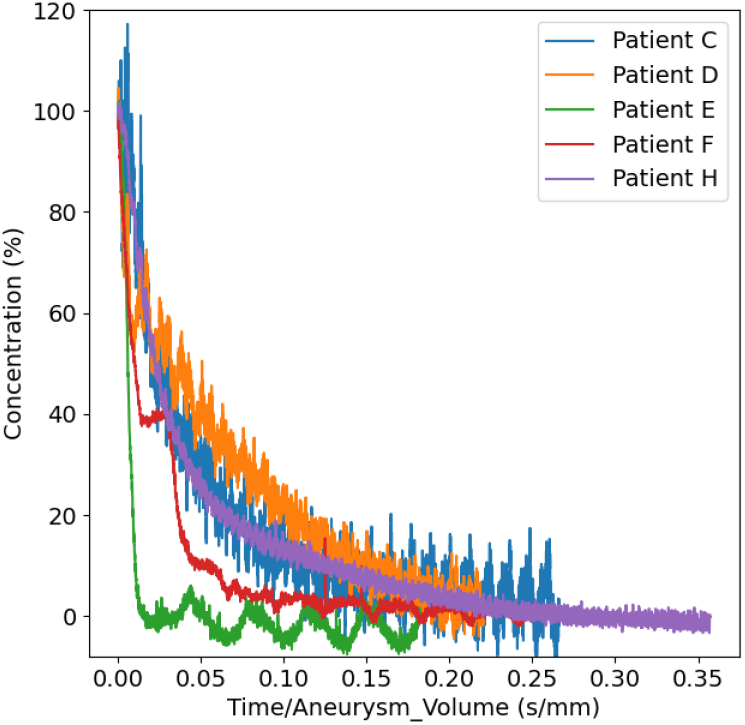
Evolution of the concentration over time normalized by the aneurysm volume for each patient. Patient G was excluded from this figure as the high variations of the concentration over time made it impossible to compare with the other patients (Figure 6G).

Table 2 presents the washout time for the six patients for three case scenarios: when the rhodamine concentration is 75%, 25%, and 0% (the total residence time). The results show that the residence time (RT) is lower for smaller volumes. However, the RT is not only defined by the size of the aneurysm, but the anatomy of the aneurysm (shape and location) seems to have an impact in RT too. Patients D and H present smaller aneurysm volumes than patients C and G respectively, but, patients D and H present larger RT than patients C and G respectively. This might be because when looking at the location of the aneurysm in Figure 2, both aneurysms (D and H) are located in bifurcation. Therefore the location and the anatomy of the aneurysm might also have an impact on the RT.

**Table 2:** Residence time where the washout is complete at c=75%, c=25% and when the washout is finished (c=0%) for the six patients. The values presented are the mean values of the three runs. Vaneurysm is the volume of each aneurysm for each patient.

| Patient | $V_{aneurysm}$ (mm<sup>3</sup>) | $T_{c=0\%}$ = RT (s) | $T_{c=25\%}$ (s) | $T_{c=75\%}$ (s) |
| --- | --- | --- | --- | --- |
| C | 93.8 | 11.16 | 4.47 | 0.948 |
| D | 90.0 | 15.09 | 6.50 | 0.29 |
| E | 32.9 | 0.43 | 0.27 | 0.14 |
| F | 38.2 | 3.88 | 1.33 | 0.148 |
| G | 413.3 | 26.67 | 13.83 | 2.48 |
| H | 279.9 | 53.77 | 13.14 | 2.98 |

#### 3.2. Comparison with coil-resolved simulations results

To compare the CFD results with the PLIF results, the evolution of concentration over time cannot be used since the boundary conditions are not the same. Indeed, the flow rate is fixed for all the patients experimentally (see section 2.2.1) whereas the simulation uses patient-specific data (see section 2.3.2). Therefore, to compare numerical and experimental results, the concentration over a pushed volume at the inlet normalized by the pore volume was analyzed. The pore volume was defined as the volume of the empty spaces in the aneurysm: the volume of the aneurysm without the volume of the coils (see Table 1). Table 3 reproduces details the different flow rates in the experiment flow rate and the simulations, with the time of the simulations and time at the end of the recording, and the total volume pushed divided by the pore volume at the end of the experiment and the simulations.

**Table 3:** Comparison between simulation and experimental results: mean flow rate, total simulation vs experimental time and volume pushed divided by the pore volume

| Patient | Experimental |  |  | Simulation |  |  |
| --- | --- | --- | --- | --- | --- | --- |
| | Mean flow rate at the inlet (mL/s) | Record washout time (s) | $V_{pushed}/V_{pore}$ at the end of the experiment | Mean flow rate at the inlet (mL/s) | Time simulation (s) | $V_{pushed}/V_{pore}$ at the end of the simulation |
| C | 3.83 | 24.99 | 1343 | 4.71 | 3.48 | 80 |
| D | 3.83 | 19.83 | 1199 | 4.27 | 3.03 | 100 |
| E | 3.83 | 5.99 | 1035 | 3.27 | 3.03 | 214 |
| F | 3.83 | 19.99 | 2816 | 4.81 | 3.03 | 264 |
| G | 3.83 | 78.80 | 920 | 2.17 | 2.68 | 17.8 |
| H | 3.83 | 99.99 | 1739 | 2.27 | 3.00 | 31.4 |

The results of the CFD analysis compared with the PLIF results are shown in Figure 9. The results of the PLIF study are shown as the mean concentration for the three runs for each patient.

**Figure 9:**
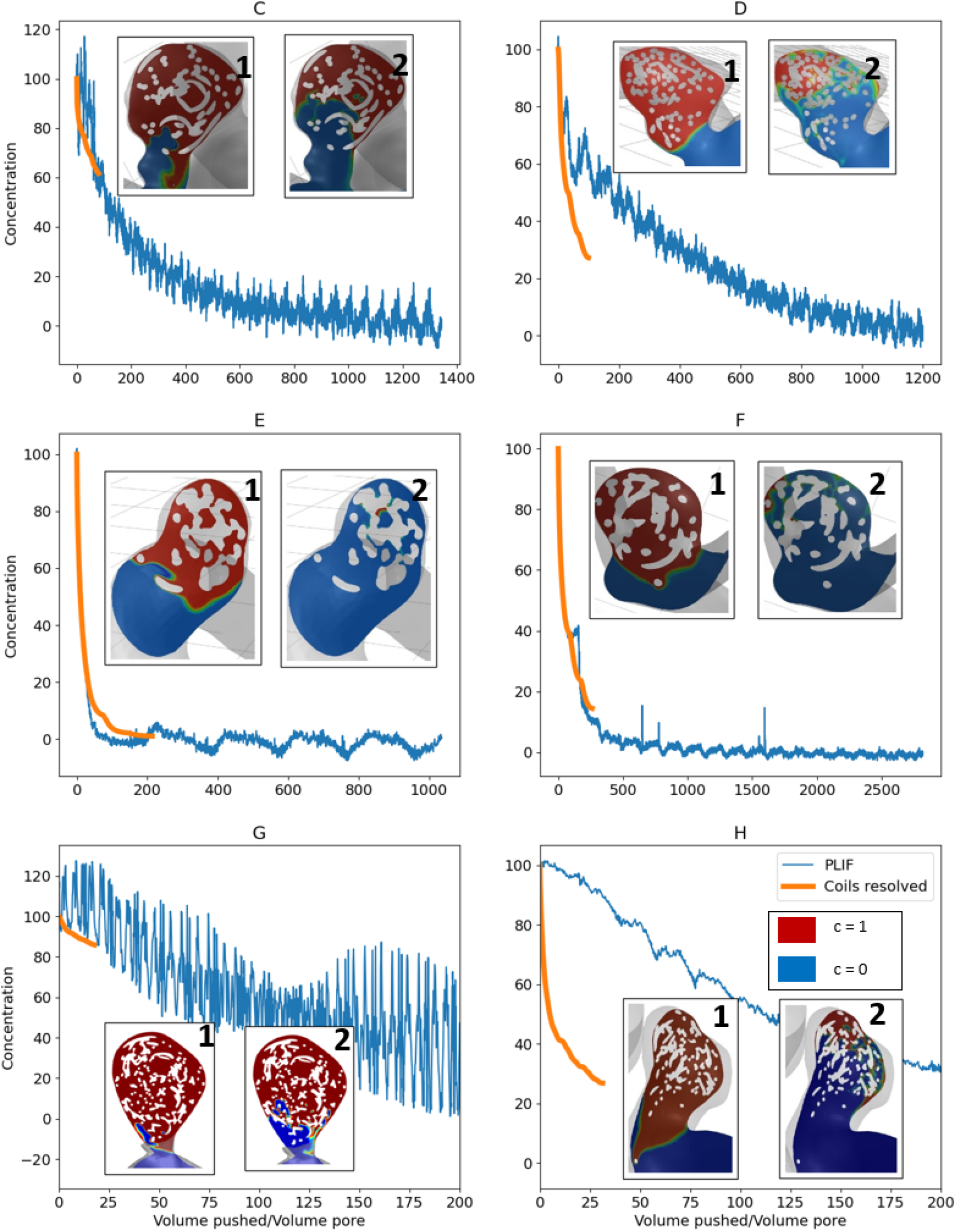
Comparison between the CFD and the PLIF study: evolution of the concentration of rhodamine B for the CFD model in orange and the experimental model in blue over pushed volume normalized by the pore volume. For each patients two captions of the evolution of concentration over time are shown: at the begining of the washout (1), and at the end of the three cardiac cycles (2).

The evolution of concentration for the simulations results show the same pattern as it did for the experimental results: the concentration decreases in the first cardiac cycle quickly and then slowly decreases until full washout. The explanation is shown by the captions of the rhodamine distribution at different instants in the simulations. At the end of the first cycle, the aneurysm neck is cleared of rhodamine, which corresponds to the areas where the velocities are higher. The caption at the end of the third cardiac cycle explains the impact of the size of the aneurysm in RT: when the aneurysms are bigger, the aneurysm presents areas with lower velocities due to the coils and the size of the aneurysm, and those areas, which are located in the aneurysm in the opposite side of the neck need more time to washout.

For patients C, E, F, and G, the numerical models reproduce the experimental results. The small differences that exist between the experimental model and the coil-resolved model might essentially due to the assumption of the coils rigidity in the numerical model. In the experimental setup, the coils are affected by the flow rate and move slightly during the cardiac cycle. This permits the movement of the rhodamine in some areas in the aneurysm where the blood velocities are very small, which impacts RT. This can be particularly perceived for patient E, where the evolution of the concentration vs the pushed volume is the same until c=20%, and then the concentration decreases slower for the coil-resolved model than for the experimental setup. Since Patient G presents a very large aneurysm volume, the RT is very high (see Table 2), which makes difficult the comparison between the experimental and the numerical work. After only three cardiac cycles the rhodamine concentration has only decrease to 80%. However, when comparing the numerical and the experimental results, they both present the same trend, therefore we can expect that the numerical model would match the experimental.

Patients D and H present a larger deviation between the experimental and the numerical results (see Figure 9). For both patients the RT is lower in the numerical than in the experimental model.

For Patient D, the origin of this deviation might be the quality of the image. As previously said, the data analysis and therefore the RT is impacted by the quality of the data and Patient D presents a poor quality image due to the arguments previously developed (see Figure 7). The origin of this problem might also be linked with the laser plane chosen to analyze the concentration over time. Patients C and D have aneurysms presenting similar volumes (93.8 and 90.0 mm^3^ respectively). When analyzing Figure 9, at the end of the third cardiac cycle, the concentration of rhodamine in the aneurysm is 60% and 25% for patients C and D respectively. These results are consistent with the captions in Figure 9. Indeed, Patient C, at the end of the third cardiac cycle is still filled with rhodamine (in red in the caption), due to its complex anatomy. However, the captions of Figure 9D show that at the end of the cardiac cycle the washout is almost complete for Patient D. This is coherent with the hemodynamic analysis: the mean velocities over one cardiac cycle in the aneurysm are 2.54 and 6.83 mm/s respectively for patients C and D. Therefore the coil-resolved model seems to be accurate. However, when analyzing Figure 8, the RT is higher for patient D than for patient C, which is not coherent with the hemodynamic results. This difference can be due to the image quality and in particular the fact that the laser plane does not illuminate a representative part of the aneurysm. The plane chosen for patient D does not illuminate the neck area as much as it does for Patient C (see Figure 7), and the plane visualized is very different from the captions in Figure 9. This particular area, the neck, is where the washout starts (see captions in Figure 9), and this area is not visualized with the camera as it is not ideally illuminated. Therefore the difference of the experimental and the numerical model might arise from the poor illumination of aneurysm D in the experimental model, and the coil-resolved model should be considered as the accurate result.

For Patient H, the origin of the deviation between the experimental and the numerical model comes from the definition of the aneurysm volume. Indeed, for this patients a part of the coils is in the parent vessel as we can see in Figure 2 and Figure 9. In the numerical model, the aneurysm is described as a envelop of the coils which includes the part of the coils in the parent vessel. This explains the quick decrease of rhodamine concentration in the numerical model, as the velocities in the parent vessel are larger that in the aneurysm and the washout is faster. In the experimental model, the aneurysm is described as the visible part in the laser plane (see Figure 7). Therefore, the only rhodamine concentration considered is visible in the aneurysm’s dome. Therefore the experiment does not match the numerical model due to the difference of the aneurysm definition. For this case, the plane did not completely illuminate the neck therefore the rhodamine in the parent vessel was visible in the camera and could not be included in the calculation of rhodamine concentration in the aneurysm.

Overall the coil-resolved simulation seems to be reproducing the same evolution of concentration observed in the experimental study, however to compare the experimental and the simulation results the laser plane needs to be adjusted so that all the regions of the aneurysm (near the neck and opposite to the neck) are used in the concentration analysis.

## 4. Discussion

This study presents an innovative experimental setup to measure RT in cerebral aneurysms treated with endovascular coiling. Studying hemodynamics experimentally in treated aneurysms presents a challenge since the coils cannot resolved using clinically available imaging studies. Most of the studies in the literature focus on measuring hemodynamics *in vivo* of the parent vessel for treated aneurysms [18]. Other studies have done PIV on idealized models treated with coils, however, the hemodynamics were only measured in the parent vessel [19]. This study presents a novel methodology for the analysis of hemodynamics in the coiled aneurysm using patient-specific anatomy and boundary conditions. In particular, the RT is an interesting parameter to be used to assess the success of the treatment. Blood coagulation is related to the blood velocity in the aneurysm, and therefore RT might be related to the coagulation and the success of the thrombus formation [20, 21, 22].

The experimental results have shown that residence time is related to the size of the aneurysm (lower for small aneurysms). Most of the areas of the aneurysm near the neck, with the highest blood velocities, get washed out in the first cardiac cycle, and then the rest of the aneurysm washes out at a slower pace. The larger the aneurysm is, the longer the washout lasts. This has been also validated numerically with the coil-resolved simulations. Other factors such as aneurysm location, morphology and parent vessel anatomy, might also be related to RT.

The coil-resolved study results are consistent with the PLIF analysis: the rhodamine concentration decreases quickly in the first cycle and then slowly decreases until full washout. The coil-resolved model allows to study at a high temporal resolution the evolution of concentration and to study the local areas where there might be stagnation and where thrombus would start to form. From the results (see Figure 9), the areas on the opposite side of the neck are where the rhodamine stays longer, and therefore where the thrombus might formed. This results are consistent with previous numerical studies where the aneurysm with coils was modeled as a porous media [23].

There are some differences between the experimental and the numerical methods linked with the boundary conditions of the model. For example, the coils are considered rigid in the numerical model whereas in the experimental setup there were small displacements. Also, the noise from the data collection of the PLIF analysis impacts this comparison.

There are some limitations related to this study. First, there is very little information that can be extracted from the PLIF analysis: only the concentration of rhodamine in the aneurysm. The analysis is done in a plane and therefore there cannot be any local analysis of rhodamine distribution. The quality of the data is very sensitive to the size of the aneurysm and to the laser plane chosen. However, it does not seem to be necessary to visualize in the camera the totality of the aneurysm volume. Patient C’s experimental results are coherent with the coil-resolved results even if only a small section of the aneurysm was illuminated (see Figure 7). However, a representative part of the aneurysm needs to be illuminated, in particular, a plane illuminating from the top of the aneurysm to the neck seems is important to study RT in a coiled aneurysm. However, this limitations could be improved by altering the optical setup: adding another laser to illuminate another part of the aneurysm or using mirrors to redirect the light of the laser.

The concentration being analyzed in the plane is also an issue for the comparison with the CFD results. We compared the mean concentration over the aneurysm volume in the coil-resolved model and the mean concentration of rhodamine in a plane. This difference might also explain the difference in some patients between the PLIF and the CFD results (example for patient D and H). Another aspect that might impact the difference between the CFD and the PLIF results is that the aneurysm volume was defined manually for the PLIF analysis. For the CFD study, the neck was defined as the convex hull envelope of the coils numerically (see section 2.1). In the PLIF analysis, the neck was defined as the coils envelop manually. The two aneurysms compared might not be exactly the same and that might impact the results of the comparison between coil-resolved and PLIF analysis.

The CFD simulations only ran for three cardiac cycles due to simulation resources. Three cardiac cycles are enough for the smaller aneurysms, and for the larger aneurysms, it can give the trend of the evolution of concentration over time. Having more computational time would improve the analysis.

Overall, this is an innovative study that includes an experimental and CFD comparison of RT in intracranial aneurysms treated with coils. The coil-resolved model used in previous studies [7, 24] was validated experimentally and can be used as a realistic model of the cerebral aneurysm treated with coils to study hemodynamics in the treated aneurysm. This analysis was done for a small sample of patients (six) and we cannot deduce from the results obtained a relation between RT and treatment success. However, this work presents a methodology to analyze, either numerically or experimentally, RT in a larger cohort of patients, and to establish, in the future, hemodynamic parameters related to treatment success.

## 5. Conclusion

Residence time was determined experimentally and numerically in six cerebral aneurysms treated with endovascular coiling. Planar laser-induced fluorescence technique was used to measure RT in phantom artery models where the aneurysms were treated with coils. The results showed that RT in the treated aneurysm is influenced by the size of the aneurysm and its anatomy. Passive scalar simulations were done in the coil-resolved model developed in previous studies to compute RT in the CFD numerical model. When comparing CFD and experimental studies, the results of the numerical simulations are consistent with the experimental results. Therefore the coil-resolved model seems to be an accurate model to measure RT. In the future, this biofidelic numerical model can be used in a larger cohort of patients to investigate the hemodynamic parameters related to treatment success.

## Acknowledgements

The 3SR lab is part of the Labex Tec 21 (Investissements d’Avenir, Grant Agreement ANR-11-LABX-0030). This work was supported by NIH/NINDS 1R01NS105692; an unrestricted educational grant from Stryker, which had no influence on the study design or results; and the generous support of the Catchot family.

## Ethical approval

Not required

## References

[1] G. Guglielmi, F. Viñuela, J. Dion, G. Duckwiler, Electrothrombosis of saccular aneurysms via endovascular approach, Journal of Neurosurgery 75 (1991) 1–7.

[2] M. Piotin, L. Spelle, C. Mounayer, M. S.-R. D. Giansante-Abud, R. Vanzin-Santos, J. Moret, Intracranial aneurysms: treatment with bare platinum coils-aneurysm packing, complex coils, and angiographic recurrence, Radiology 243 (2) (2007) 500–8.

[3] E. Crobeddu, G. Lanzino, D. Kallmes, H. Cloft, Review of 2 decades of aneurysm-recurrence literature, part 1: reducing recurrence after endovascular coiling, AJNR Am J Neuroradiol 34 (2) (2013) 266–70.

[4] S. Tamatani, Y. Ito, H. A. T. K. S. Takeuchi, R. Tanaka, Evaluation of the stability of aneurysms after embolization using detachable coils: correlation between stability of aneurysms and embolized volume of aneurysms, AJNR Am J Neuroradiol 23 (5) (2002) 762–7.

[5] H. Meng, Z. Wang, Y. Hoi, L. Gao, E. Metaxa, D. Swartz, J. Kolega, Complex hemodynamics at the apex of an arterial bifurcation induces vascular remodeling resembling cerebral aneurysm initiation, Stroke 38 (6) (2007) 1924–31.

[6] R. J. Damiano, D. Ma, J. Xiang, A. H. Siddiqui, K. V. Snyder, H. Meng, Finite element modeling of endovascular coiling and flow diversion enables hemodynamic prediction of complex treatment strategies for intracranial aneurysm, Journal of Biomechanics 48 (12) (2015) 3332–3340.

[7] M. R. Levitt, M. C. Barbour, S. R. du Roscoat, C. Geindreau, V. K. Chivukula, P. M. McGah, J. D. Nerva, R. P. Morton, L. J. Kim, A. Aliseda, Computational fluid dynamics of cerebral aneurysm coiling using high-resolution and high-energy synchrotron X-ray microtomography: comparison with the homogeneous porous medium approach, Journal of NeuroInterventional Surgery 0 (2016) 1–6.

[8] B. Seibert, R. Tumala, R. Chow, A. Farida, S. Mousavi, A. Divani, Intracranial aneurysms: review of current treatment options and outcomes, Front Neurol. 2011 (12) (2011) 2:45.

[9] M. Barbour, P. McGah, C. Ng, A. Clark, K. Gow, A. Aliseda, Convective Leakage Makes Heparin Locking of Central Venous Catheters Ineffective Within Seconds: Experimental Measurements in a Model Superior Vena Cava, ASAIO J. 61 (6) (2015) 701–9.

[10] M. Barbour, K. Gow, A. Aliseda, Dominated Heparin Leakage From Multiple Catheter Designs: An In Vitro Experimental Study, ASAIO J. 64 (5) (2015) 94–104.

[11] K. Venkat, M. Levitt, A. Clark, S. R. du Roscoat, L. Kim, A. Aliseda, Reconstructing patient-specific cerebral aneurysm vasculature for in vitro investigations and treatment efficacy assessments, Journal of Clinical Neuroscience 61 (2019) 153–59.

[12] J. Schindelin, I. Arganda-Carreras, E. Frise, V. Kaynig, M. Longair, T. Pietzsch, A. Cardona, Fiji: an open-source platform for biological-image analysis, Nature Methods 9 (7) (2012) 676–82.

[13] J. Wilm, Iterative closest point, matlab central file exchange (2020). URL https://www.mathworks.com/matlabcentral/fileexchange/27804-iterative-closest-point

[14] M. Ford, N. Alperin, S. Lee, D. Holdsworth, D. Steinman, Characterization of volumetric flow rate waveforms in the normal internal carotid and vertebral arteries, Physiol Meas 26 (4) (2005) 477–88.

[15] K. Chandran, S. Rittgers, A. Yoganathan, Biofluid mechanics: the human circulation. Second edition., 2012.

[16] P. McGah, D. Leotta, K. Beach, J. Riley, A. Aliseda, A longitudinal study of remodeling in a revised peripheral artery bypass graft using 3D ultrasound imaging and computational hemodynamics, Journal of biomechanical engineering 133 (4) (2011) 041008.

[17] M. Levitt, P. McGah, A. Aliseda, P. Mourad, J. Nerva, S. Vaidya, R. Morton, B. Ghodke, L. Kim, Cerebral aneurysms treated with flow-diverting stents: computational models with intravascular blood flow measurements, AJNR. American journal of neuroradiology 35 (1) (2014) 143–148.

[18] M. R. Levitt, S. S. Vaidya, J. C. Mai, D. K. Hallam, L. J. Kim, B. V. Ghodke, Balloon Test Occlusion with the Doppler Velocity Guidewire, Journal of Stroke and Cerebrovascular Diseases 21 (8) (2012) 909.e1–909.e4.

[19] P. Nair, B. Chong, A. Indahlastari, J. Ryan, C. Workman, M. H. Babiker, H. Y. F. Hooman, C. Baccin, D. Frakes, Hemodynamic Characterization of Geometric Cerebral Aneurysm Templates Treated With Embolic Coils, Journal of Biomechanical Engineering 138 (2).

[20] J. Jesty, W. Yin, P. Perrotta, D. Bluestein, Platelet activation in a circulating flow loop: combined effects of shear stress and exposure time, Platelets 14 (3) (2003) 143:9.

[21] J. Cao, S. Rittgers, Particle motion within in vitro models of stenosed internal carotid and left anterior descending coronary arteries, Ann Biomed Eng. 26 (2) (1998) 190:9.

[22] A. Gear, Platelet adhesion, shape change, and aggregation: rapid initiation and signal transduction events, Can J Physiol Pharmacol 72 (3) (1994) 285:94.

[23] A. Mitsos, N. Kakalis, Y. Ventiko, J. Byrne, Haemodynamic simulation of aneurysm coiling in an anatomically accurate computational fluid dynamics model: technical note, Neuroradiology 50 (4) (2008) 341–347.

[24] M. Barbour, Computational and Experimental Investigation into the Hemodynamics of Endovascularly Treated Cerebral Aneurysms, 2018.

